# A single K^+^-binding site in the crystal structure of the gastric proton pump

**DOI:** 10.1101/608851

**Authors:** Kenta Yamamoto, Vikas Dubey, Katsumasa Irie, Hanayo Nakanishi, Himanshu Khandelia, Yoshinori Fujiyoshi, Kazuhiro Abe

## Abstract

The gastric proton pump (H^+^,K^+^-ATPase), a P-type ATPase responsible for gastric acidification, mediates electro-neutral exchange of H^+^ and K^+^ coupled with ATP hydrolysis, but with an as yet undetermined transport stoichiometry. Here we show crystal structures at a resolution of 2.5 Å of the pump in the E2-P transition state, in which the counter-transporting cation is occluded. We found a single K^+^ bound to the cation-binding site of H^+^,K^+^-ATPase, indicating an exchange of 1H^+^/1K^+^ per hydrolysis of one ATP molecule. This fulfils the energy requirement for the generation of a six pH unit gradient across the membrane. The structural basis of K^+^recognition is resolved, supported by molecular dynamics simulations, and this establishes how H^+^,K^+^-ATPase overcomes the energetic challenge to generate an H^+^ gradient of more than a million-fold – the highest cation gradient known in any mammalian tissue – across the membrane.

## Introduction

While the closely-related Na^+^,K^+^-ATPase mediates electrogenic transport of three Na^+^ and two K^+^ ions coupled with the hydrolysis of one ATP molecule (*1*), the number of transported cations in the electroneutral operation (*2,3*) of the gastric H^+^,K^+^-ATPase remains unclear. Under physiological conditions, intact parietal cells with an internal pH of approximately seven must generate a pH gradient of at least six pH units (*4*). The secretion of a relatively voluminous flow of gastric acid requires the expenditure of considerable cellular energy. Taking reasonable values of pH 7 and 120 mM K^+^ for the intracellular condition with measured pH (1.0) and K^+^ concentration (10 mM) of the gastric juice gives concentration gradients across the membrane of 10^6^ and 12 times for H^+^ and K^+^, respectively. The sum of chemical potentials is about −10 kcal/mol. The reported free energy derived from ATP hydrolysis in the parietal cell is about −13 kcal/mol (*5*). Therefore, the ratio of H^+^ transported to ATP hydrolyzed must be approximately 1, and cannot be as large as 2, when the gastric pH approaches 1 (Figure 1). A previous investigation of H^+^ transport shows stoichiometric H^+^/ATP hydrolysis at pH 6.1-6.9 (*6*), consistent with the above. However, this cannot be the case when luminal pH is neutral to weakly acidic (Figure 1). A separate investigation, on the other hand, showed two H^+^ transported per one ATP hydrolyzed at pH 6.1 (*7*). It was speculated that the number of H^+^ transported may change from two (at neutral pH) to one as the luminal pH decreases (*8,9*). It remains unclear if the transport of two H^+^ is physically possible at neutral luminal pH. The best method of resolving the disputed transport stoichiometry of the pump is by direct observation of the number of counter-transported cations occluded in the H^+^,K^+^-ATPase at neutral pH, given that the transport is approximately electroneutral (*2,3*).

**Figure 1.**
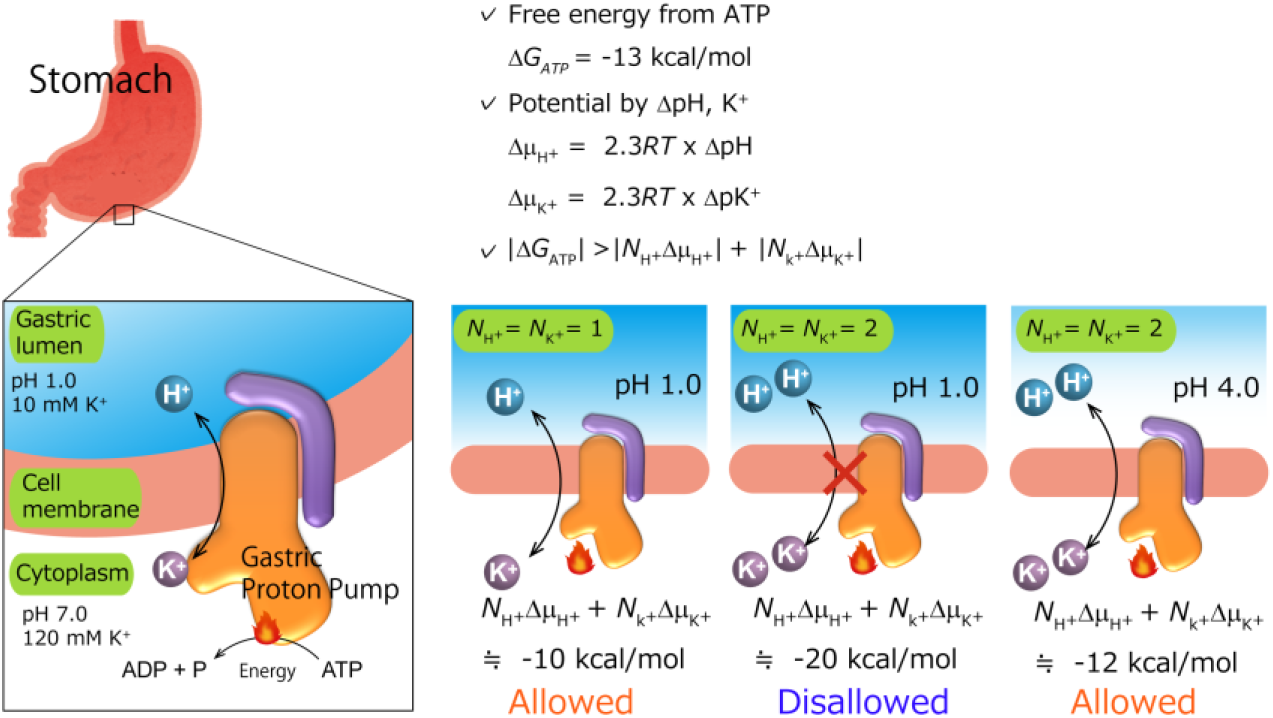
Transport stoichiometry and free energy from ATP hydrolysis. The free energy derived from ATP hydrolysis, *ΔG_ATP_*, calculated from *ΔG*’_0_ and the measured intracellular concentrations of ATP, ADP and Pi in the parietal cell is about −13 kcal/mol (*5*) as described in the previous report. Under physiological conditions in intact parietal cells with an internal pH of approximately 7, a pH gradient of at least 6 pH units must be created. The maximum electrochemical gradient, *Δμ_i_*, that can be formed by an ion-transporting ATPase is a function of the free energy of ATP hydrolysis. Taking reasonable values of pH 7 and 120 mM K^+^ for intracellular conditions with the measured pH (1.0) and K^+^ concentration (10 mM) of the gastric juice gives concentration gradients across the parietal cell membrane of 10^6^ and 12 times for H^+^ and K^+^, respectively. For an electro-neutral H^+^/K^+^ exchange pump where *N*_*H*+_ = *N*_*K*+_ = 1, the sum of chemical potentials is about −10 kcal/mol, and is within the range of ATP free energy (*6*). However, if the exchange of two cations is assumed where *N*_*H*+_ = *N*_*K*+_ = 2, the reaction is thermodynamically disallowed at pH 1. Therefore, the ratio of H^+^ transported to ATP hydrolyzed must be approximately 1, and cannot be as large as 2, when gastric pH is around 1. However, this cannot be the case when luminal pH is neutral to weakly acidic, *e.g*., at pH 4. A different postulate is that two H^+^ are transported per ATP molecule hydrolyzed under these conditions (*7*), and that the number of transported H^+^, and therefore K^+^ as well, changes from 2 to 1 as luminal pH decreases.

## Results

To this end, we determined crystal structures at neutral pH, in order to determine the maximum capacity of K^+^ occlusion. According to the transport cycle of H^+^,K^+^-ATPase (*10*) (Figure S1), binding of counter-transporting K^+^ induces luminal gate closure and accelerates dephosphorylation of the auto-phosphorylated intermediate, E2P. Subsequently, the enzyme moves to the E2-P transition state in which counter-transported K^+^ is occluded (*9*). For crystallization, the transition state phosphate analogs (*11*) (magnesium fluoride (MgF_4_^2-^) or aluminum fluoride (AlF_4_^-^)) with counter-transporting cation (K^+^ or its congener Rb^+^) were applied (Figure S1), respectively, to the wild-type (WT) H^+^,K^+^-ATPase. However, crystals diffract poorly in these conditions, and the resolution was limited to 4.2 Å - insufficient for the precise definition of the K^+^-coordination. Better-resolving crystals were obtained by using the Tyr799Trp mutant on the α-subunit (YW). Tyr799 is located at the entrance of the luminal-facing cation gate, 12 Å distant from the K^+^ binding site, and its mutation does not affect K^+^ binding site directly. To our surprise, however, the Tyr799Trp mutant shows the highest ATPase activity in the absence of K^+^ and the activity decreases with increasing K^+^ concentration, in marked contrast to the K^+^-dependent increase in ATPase activity of the wild-type enzyme (Figure S1). The observed K^+^-independent ATPase activity of Tyr799Trp in the absence of K^+^ indicates that the luminal gate of this mutant is spontaneously closed, like previously reported constitutive-active mutants (*12*), *i.e*., the Tyr799Trp mutation exerts a molecular signal that induces E2P dephosphorylation when K^+^ is bound to the cation binding site. Decreasing ATPase activity of Tyr799Trp with increasing K^+^ concentration (*K*_0.5,cyto_ = ~5 mM) can be interpreted as being due to K^+^-occlusion from the cytoplasmic side, which is also observed in the wild-type enzyme albeit with much lower affinity (*K*_0.5,cyto_ = ~200 mM, *13*). Furthermore, in the presence of an inhibiting concentration of the K^+^-competitive blocker vonoprazan, Tyr799Trp shows high-affinity K^+^-activation of ATPase activity. These data indicate that, despite the unique properties of luminal gate closure, the cation-binding site of Tyr799Trp is intact, and capable of high-affinity K^+^-binding. The thermal stability (*14*) of Tyr799Trp is significantly higher in the presence of K^+^ and MgF_4_^2-^ than MgF_4_^2-^ alone (Figure S1), qualitatively indicating K^+^-occlusion in the Tyr799Trp mutant. This K^+^-occluded E2-P transition state is the most stable conformation amongst all evaluated conditions for Tyr799Trp, in contrast to the WT enzyme in which inhibitor-bound and luminal-open E2BeF is the most stable. We therefore conclude that Tyr799Trp prefers the luminal-closed K^+^-occluded state, (K^+^)E2-P, and is thus suitable for the structural analysis of the K^+^-occluded form.

As expected, crystals were significantly improved using the Tyr799Trp mutant, and provided a 2.5 Å resolution structure in the best case [YW(K^+^)E2-MgF_4_^2-^] (Figure 2). We analyzed several crystal structures in the presence of different combinations of K^+^ or Rb^+^, and AlF_4_^-^ or MgF_4_^2-^, all of which mimic the K^+^-occluded E2-P transition state and are indistinguishable in molecular conformation (Figure S2). Although the analyzed resolution is limited, the structure of the wild-type enzyme also shows almost the same molecular conformation as that of the Tyr799Trp mutant (Figure S2). We therefore use YW(K^+^)E2-MgF_4_^2-^ structure analyzed at the best resolution in the following discussion (Table S1). The overall structure of H^+^,K^+^-ATPase YW(K^+^)E2-MgF_4_^2-^ (Figure 2A) is very close to the corresponding structure of Na^+^,K^+^-ATPase (*15,16*) (Figure S2). However, instead of the two K^+^ ions occluded in the transmembrane cation-binding site of the Na^+^,K^+^-ATPase, only a single K^+^ is observed in that of H^+^,K^+^-ATPase (Figure 2). The observed single K^+^-binding is confirmed by the anomalous difference Fourier maps of YW(Rb^+^)E2-MgF_4_^2-^ (Figure 2B), YW(Rb^+^)E2-AlF_4_^-^ and YW(K^+^)E2-MgF_4_^2-^ structures (Figure S2), unambiguously showing a single strong peak at the cation binding site located between TM4, 5 and 6 in the middle section of the membrane. The presence of saturating concentrations of cation (400 mM KCl, or RbCl) in the crystallization buffer ensures high occupancy of K^+^ at the cation-binding site, although several other binding sites, presumably low-affinity and/or non-specific, were determined in the cytoplasmic domains (Figure S2). A single anomalous peak at the cation binding site is also found in the wild-type WT(Rb^+^)E2-MgF_4_^2-^ state (Figure S2), thus excluding the possibility that the observed single K^+^-binding is due to the artifact of Tyr799Trp mutation.

**Figure 2.**
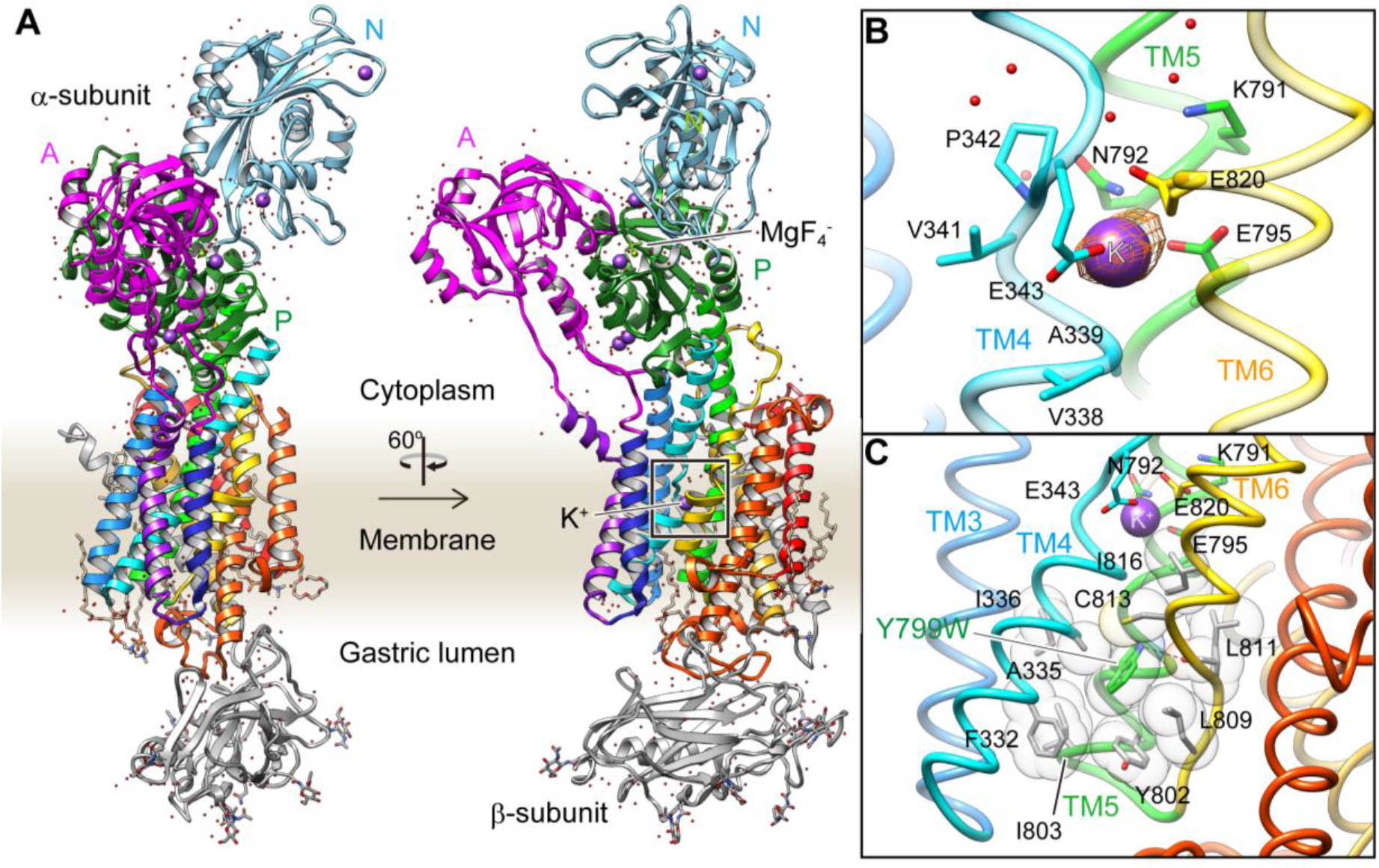
Crystal structure of the K^+^-occluded E2-P transition state of H^+^,K^+^-ATPase. A, Overall structure of the K^+^-occluded E2-MgF_4_^2-^ state [YW(K^+^)E2-MgF_4_^2-^] in ribbon representations. For the α-subunit, the three cytoplasmic domains (A, P, N) are shown in different colors. Color of the TM helices gradually changes from purple to red (TM1-TM10). The β-subunit with a single TM helix and six *N*-glycosylation sites in the ecto-domain are shown in grey. Phospholipids, a cholesterol and detergent molecules are also modeled (sticks). Red dots and purple spheres represent water molecules and K^+^ ions, respectively. **B**, TM K^+^-binding site viewed from parallel to membrane plane. Orange mesh represents anomalous density map of Rb^+^-bound form [(Rb^+^)E2-MgF_4_^2-^] with 8 σ contour level. Amino acids that contribute to the K^+^-coordination are shown in sticks. **C**, The hydrophobic gate centered around Tyr799Trp (green), with surrounding hydrophobic residues (grey) are shown. Dotted line indicates a hydrogen bond between a nitrogen atom of Trp residue and a main chain oxygen of L811.

Crystals were generated at the close to neutral pH of 6.5, the same condition used in previous *in vitro* H^+^ transport measurements (*6,7*). Assuming electro-neutral transport in the H^+^,K^+^-ATPase (*2,3*), if two H^+^’s are transported at neutral pH, as suggested by a previous *in vitro* measurement (*7*), then two K^+^ ions must be counter-transported. A single bound K^+^ observed in the crystal structure therefore strongly indicates that H^+^,K^+^-ATPase transports only one K^+^ ion at once, and therefore only one H^+^ in the opposite direction, for every ATP hydrolyzed at neutral pH. Single K^+^-binding in H^+^,K^+^-ATPase is also supported by a Hill coefficient for K^+^ of close to 1.0 (Figure S1), in marked contrast to a Hill coefficient of 1.5 for the K^+^-dependence of Na^+^,K^+^-ATPase (*17*) in which two K^+^ ions are occluded at the cation-binding site.

Different from the previously reported Rb^+^-bound and luminal-open (SCH)E2BeF structure of H^+^,K^+^-ATPase (*12*), the cation-binding site is now tightly shielded from the luminal solution in the YW(K^+^)E2-MgF_4_^2-^ state, by an extensive hydrophobic cluster centered around Tyr799Trp (Figure 2C) at the luminal gate. Substitution of Tyr799 to Trp enhances hydrophobic interactions with surrounding amino acids, the interaction of which may be strong enough for spontaneous luminal gate closure (*12*) as suggested by the highest ATPase activity of the Tyr799Trp mutant in the absence of K^+^ (Figure S1, Figure S3). A hydrogen bond between the nitrogen of the Trp799 residue and main chain carbonyl oxygen of Leu811 (3.3 Å, Figure 2C, Figure S3) provides for the positioning of a Trp rotamer suitable for the hydrophobic interactions with its surroundings, and stabilizes its conformation. Molecular dynamics (MD) simulations confirm the presence of this hydrogen bond, and also show that in the WT enzyme, the hydroxyl group of Tyr799 makes a similar hydrogen bond with the main chain carbonyl oxygen of Leu811 (3.0 Å, Figure S3, Movie S1). However, the Tyr side chain is smaller than that of Trp, and thus forms a weaker hydrophobic network with the surrounding residues. Although the Leu811Pro mutant is unfolded, the Leu811Gly mutant in the background of Tyr799Trp shows weak K^+^-dependence in its ATPase activity (Figure S3), suggesting that glycine substitution might alter the main-chain trace near Leu811, thus weakening the hydrogen-bond interaction with Trp799. Hydrophilic serine substitution of the surrounding hydrophobic residues in the Try799Trp background does not significantly change the inverse K^+^-dependence profile of Tyr799Trp activity, except for Tyr799Trp + Ile803Ser (YWIS) which shows a K^+^-dependent increase in its ATPase activity. However, there still remains a K^+^-independent ATPase fraction in the absence of K^+^. In the background of YWIS, an additional third mutation (Leu809Ser, Cys813Ser or Ile816Ser) restores K^+^-dependent ATPase activity approximating that of the wild-type enzyme activity. These data strongly suggest that spontaneous gate closure of the Tyr799Trp mutant is caused by extensive hydrophobic interactions with its surroundings, and these are facilitated by the favorable rotamer position of Tyr799Trp guided by the hydrogen-bond between Trp799 and Leu811 main chain. Wild-type-like K^+^-dependence can be restored by additional mutagenesis based on the observed structure of YW(K^+^)E2-P. We therefore conclude that the luminal-closed molecular conformation spontaneously induced by the Try799Trp mutation is not an artifact, and the driving force for the gate closure is essentially the same as in the wild-type enzyme.

Comparison of the luminal-open E2P ground state [(von)E2BeF structure with bound vonoprazan, a specific inhibitor for H^+^,K^+^-ATPase (*12*)] and K^+^-occluded and luminal-closed E2-P transition state [YW(K^+^)E2-MgF_4_^2-^ structure] reveals several key conformational rearrangements upon luminal gate closure (Figure S4). Gate closure brings the luminal portion of TM4 (TM4L) close to TM6, effectively capping the cation-binding site from the luminal side of the membrane. This lateral shift of TM4L is coupled to the vertical movement of the TM1,2 helix bundle connected to the A domain, leading to the ~30° rotation of the A domain that induces dephosphorylation of the aspartylphosphate. The molecular events required for the luminal gate closure have been extensively studied in SERCA (*18–20*), and the same mechanism is observed in the H^+^,K^+^-ATPase, confirming low-resolution maps of electron crystallography (*21,22*). The lateral shift of TM4L not only blocks the physical path of the cation from the luminal solution, but also brings main chain oxygen atoms important for the high-affinity K^+^-coordination to their optimal positions, as described later.

Our crystal structure defines a high-affinity K^+^-binding site of H^+^,K^+^-ATPase (Figure 3, Movie S2), with the coordination geometry of K^+^ by the surrounding amino acids evident at 2.5 Å resolution. The bound single K^+^ in H^+^,K^+^-ATPase is located at a position corresponding to site II of Na^+^,K^+^-ATPase (2K^+^)E2-MgF state (*15*). The bound K^+^ is coordinated by eight oxygen atoms located within 4 Å distance (Table 1). Of these, five make a large contribution to K^+^ coordination (within 3 Å), they include three oxygen atoms from main-chain carbonyls (Val338, Ala339 and Val341) and two from side chain carboxyl groups (Glu343 and Glu795). The total valence (*23,24*) for the bound K^+^ is 1.05 (optimal valence for K^+^ is 1.00), indicating K^+^ is almost ideally coordinated by the surrounding oxygen atoms (Table 1). Similar results were obtained for Rb^+^ coordination in the YW(Rb^+^)E2-MgF_4_^2-^ (valence 1.14) and YW(Rb^+^)E2-AlF_4_^-^ (valence 1.07) structures, but clearly different from the previously reported Rb^+^-bound (SCH)E2BeF structure (*12*) (valence 0.39, see Table1). As the TM4 helix is unwound at Pro342, coordination by the main chain carbonyl groups becomes possible (Figure 3). These carbonyls, in addition to the Glu343 carboxyl, likely determine the positioning of TM4L upon luminal gate closure coupled with K^+^-occlusion (Figure S4). It seems to be a natural consequence that these carbonyls dominate K^+^ coordination, as is found in Na^+^,K^+^-ATPase (*15*) and K^+^-channels (*25*). Unwinding of TM4 also provides a negative dipole moment (δ^-^) at the K^+^ site, which is located on the extension of TM4L at the helix breaking point.

**Figure 3.**
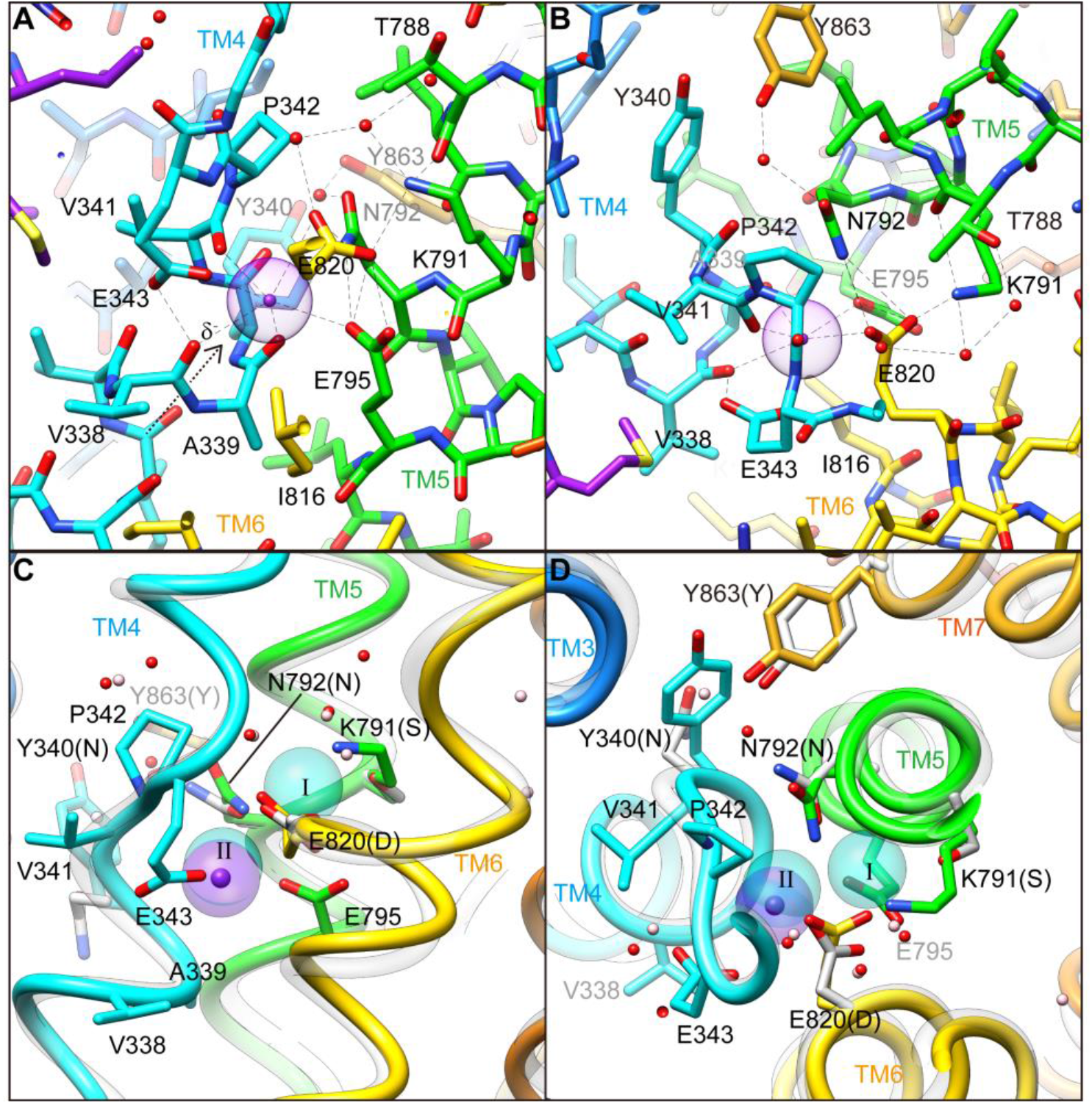
K^+^-binding site. (**A** and **B**) Close-up of the transmembrane cation-binding site in H^+^,K^+^-ATPase YW(K^+^)E2-MgF_4_^2-^ state in stick representation, viewed approximately parallel to the membrane from TM6 side (**A**) or approximately perpendicular to the membrane from cytoplasmic side (**B**). Dotted lines indicate atoms within 3.5 Å distance of neighboring atoms, presumably making hydrogen bonds or electrostatic interaction. A purple dot indicates bound K^+^ with its Stokes radius shown as a transparent sphere. Water molecules (red dots) are also indicated. (**C** and **D**) (2K^+^)E2-MgF_4_^2-^ state of Na^+^,K^+^-ATPase (*15*) (grey ribbons) is superimposed on the corresponding reaction state of H^+^,K^+^-ATPase YW(K^+^)E2-MgF_4_^2-^ (color ribbons), viewed from membrane (C) or cytoplasmic side (**D**). For clarity, amino acid residues contributing to the K^+^ coordination in H^+^,K^+^-ATPase are indicated, with their corresponding amino acids in Na^+^,K^+^-ATPase in parentheses. Residues of Asn331, Ser782, Asn783, Asp811, Tyr854 in Na^+^,K^+^-ATPase, corresponding to Tyr340, Lys791, Asn792, Glu820 and Tyr863 in H^+^,K^+^-ATPase, respectively, are shown because three of these amino acids are not conserved, others are conserved but different conformation in the structure. Blue spheres indicate positions of K^+^ binding site I and II, and pink dots are water molecules in Na^+^,K^+^-ATPase structure.

**Table 1.** Coordination geometry and partial valence in the K^+^-binding site of H^+^,K^+^-ATPase in (K^+^)E2-P analogue states. Only oxygen atoms within 4 Å from K^+^ are included for the valence calculation. Partial valence is calculated for K^+^ and Rb^+^ in each corresponding crystal structures.

| Structure | Rb<sup>+</sup>-(SCH)E2BeF |  | YW(K<sup>+</sup>)E2-MgF |  | YW(Rb<sup>+</sup>)E2-MgF |  | YW(Rb<sup>+</sup>)E2-AlF |  |
| --- | --- | --- | --- | --- | --- | --- | --- | --- |
| Amino acids | Distance, Å | Valence | Distance, Å | Valence | Distance, Å | Valence | Distance, Å | Valence |
| V338 O | 4.57 | - | 2.90 | 0.11 | 3.00 | 0.12 | 3.09 | 0.1 |
| A339 O | 3.23 | 0.07 | 2.62 | 0.28 | 2.68 | 0.27 | 2.63 | 0.31 |
| V341 O | 2.89 | 0.16 | 2.56 | 0.34 | 2.56 | 0.37 | 2.63 | 0.31 |
| E343 Oε1 | 3.41 | 0.05 | 2.93 | 0.10 | 2.96 | 0.13 | 3.07 | 0.10 |
| E343 Oε2 | 3.42 | 0.05 | 3.74 | 0.01 | 3.63 | 0.03 | 3.72 | 0.03 |
| E795 Oε1 | 4.05 | - | 2.77 | 0.17 | 2.99 | 0.12 | 2.88 | 0.16 |
| E820 Oε1 | 3.56 | 0.04 | 3.32 | 0.03 | 3.21 | 0.08 | 3.38 | 0.05 |
| E820 Oε2 | 3.68 | 0.03 | 3.90 | 0.01 | 3.98 | 0.02 | 3.95 | 0.02 |
| Total valence |  | 0.39 |  | 1.05 |  | 1.14 |  | 1.07 |

Side chain oxygens from Glu343 and Glu795 also contribute to K^+^ coordination. However, even in the presence of a positive charge (K^+^ or Rb^+^), the Glu343 side chain is not attracted to the bound cation. The charge-conserved mutant of Glu343Asp shows no detectable ATPase activity due to its inability to induce K^+^-dependent dephosphorylation (*26*), indicating that having a negative charge at this position is not essential for the K^+^-coordination. Glu343Gln reduces K^+^-affinity (*12*), but to a far lesser extent than Glu343Asp, suggesting that Glu343 is likely protonated in the crystal structure. The short distance between the Glu343 carboxyl and Val341 carbonyl (2.6 Å) suggest protonation of Glu343. The juxtaposition of Glu795 and Glu820 indicates that at least one of the carboxyls must be protonated (*12,27*). Based on the almost identical K^+^-affinity of WT and the charge-neutralized mutant Glu795Gln, Glu795 is the one expected to be protonated (*12*). Therefore, compared with other coordinating side chain oxygens, the close location (2.8 Å) of Glu795, and thus larger contribution to K^+^-coordination is not due to a negative charge, but to an appropriate size of the Glu side chain, as can be deduced from the reduction in either ATPase activity and K^+^-affinity of the Glu795Asp mutant (*12*). The relatively distant location of Glu820 from the bound K^+^ is due to a salt bridge with Lys791. This counteracts the negative charge at the Glu820 side chain, which is essential for the H^+^ extrusion to the luminal solution in the previous step of the transport cycle, namely, the luminal-open E2P state (*12*).

In the cation-binding site of the H^+^,K^+^-ATPase structure, there is not enough space to accommodate a second K^+^ at the position corresponding to the site I of Na^+^,K^+^-ATPase (Figure 3C,D, Figure 4). This restricted structure is due to the position of the Lys791 side chain and that of its salt bridge partner Glu820. These two amino acids are invariant for the gastric α1 isoform of H^+^,K^+^-ATPase, but are replaced with Ser782 and Asp811 (shark α1 sequence), respectively, in Na^+^,K^+^-ATPase (Figure S5). In particular Ser782 in Na^+^,K^+^-ATPase plays an important role in K^+^-accommodation at site I (*15,16*). Replacing Ser with the bulky and positively-charged Lys791 in H^+^,K^+^-ATPase, would prevent K^+^ binding at a site I position. In fact, when the two structures (H^+^,K^+^-ATPase and Na^+^,K^+^-ATPase) were superimposed, K^+^ at site I and a coordinating water molecule in Na^+^,K^+^-ATPase sterically clashed with the H^+^,K^+^-ATPase Lys791 side chain (Figure 4), the position of which is defined by its salt-bridge partner Glu820 and a hydrogen bond with the main chain carbonyl of Thr788 (Figure 3). In contrast to the important contribution of the side chain carbonyl of Asn783 for K^+^ coordination in Na^+^,K^+^-ATPase (*15*), the carbonyl oxygen of the Asn792 side chain in H^+^,K^+^-ATPase is facing towards the opposite side of K^+^, and the Asn792 amino group stabilizes the positions of Glu795 and Glu820 by making hydrogen bonds (Figure 3C,D). In the Na^+^,K^+^-ATPase structure, Asn783 is stabilized by forming a hydrogen bond with Tyr854 in TM7 (*15*). In H^+^,K^+^-ATPase, because of Tyr340 (corresponds to Asn331 in Na^+^,K^+^-ATPase) in TM4, the side chain of Tyr863 (TM7, corresponds to Tyr854 in Na^+^,K^+^-ATPase) is removed from the position observed in Na^+^,K^+^-ATPase, which brings a water molecule in between Asn792 and Tyr863 in H^+^,K^+^-ATPase, and stabilizes the Asn792 side chain. Except for the observed differences described above, the cation-binding site of these two related ATPases is surprisingly similar, not only with respect to the positions of the side chains, but also with those of water molecules.

**Figure 4.**
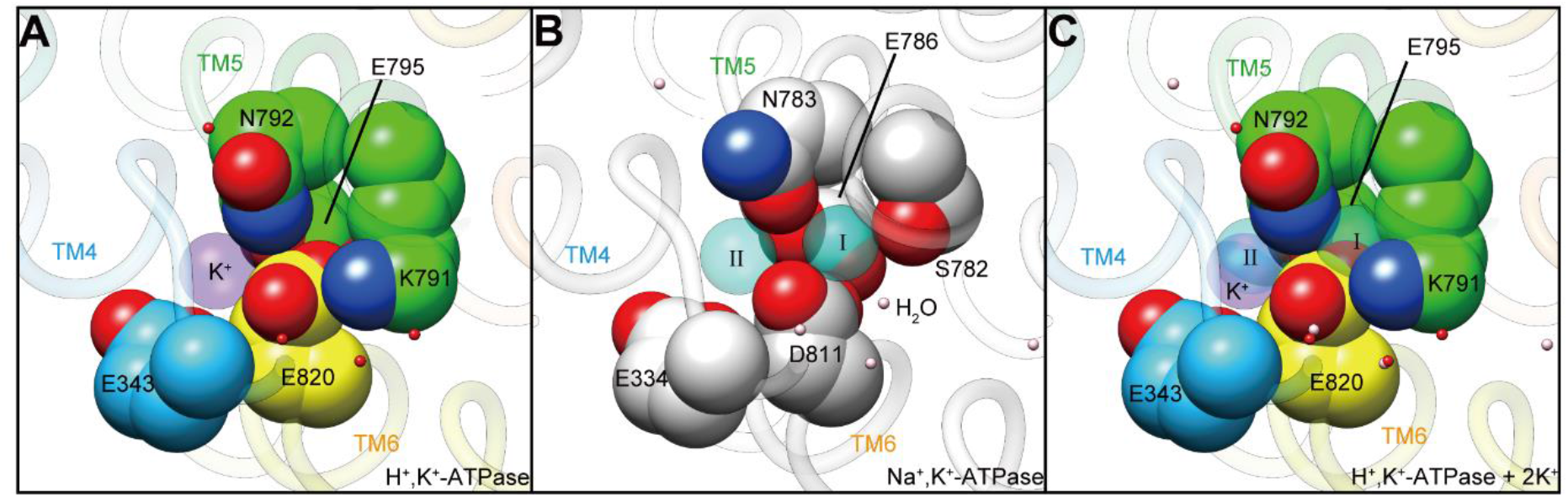
Comparison of the cation-binding site between H^+^,K^+^- and Na^+^,K^+^-ATPase. Cation-binding sites of H^+^,K^+^-ATPase YW(K^+^)E2-MgF_4_^2-^ (**A**) and Na^+^,K^+^-ATPase (2K^+^)E2-MgF_4_^2-^ (**B**) are shown. Two K^+^ ions occluded in Na^+^,K^+^-ATPase (I, II) were superimposed on the H^+^,K^+^-ATPase structure (**C**), showing significant steric clash between K^+^ at site I, and Lys791 and Glu820. Only side chains that is important for the K^+^ coordination are displayed as space-fill models. Water molecules in H^+^,K^+^-ATPase and Na^+^,K^+^-ATPase are shown as red and pink dot, respectively. Figures are viewed from approximately perpendicular to the membrane from the cytoplasmic side. Color codes as in Fig. 2.

The protonation status of acidic residues cannot be determined at 2.5 Å resolution, and therefore we turned to MD simulations with different protonation combinations of the acidic side chains at the cation-binding site (Figure 5). Besides the K^+^-coordinating three glutamates (Glu343, Glu795 and Glu820), there are three other acidic amino acids near the cation-binding site (Figure S6), which are also considered in the system. Two acidic residues Asp824 (TM6) and Glu936 (TM8), located on the opposite side of the Glu795-Glu820 pair with Lys791 in between, are juxtaposed (2.8 Å), and therefore one of these acidic side chains is expected to be protonated. Indeed, p*K*_a_ distributions obtained from MD simulations always suggest deprotonation of Asp824 and protonation of Glu936, irrespective of the protonation state of other acidic residues in the cation-binding site. When protonation is assumed for Asp824, both Asp824 and Glu936 show bimodal *p*K_a_ distributions (Figure S6), which is unexpected for the stable, occluded cation-bound conformation of the H^+^,K^+^-ATPase. Although the Asp824-Glu936 pair is rather isolated from the K^+^-binding site in the E2-P form of H^+^,K^+^-ATPase, this acidic residue pair seems to be capable of neutralizing the positive charge of Lys791 when it flips – as is expected in the E1 conformation. The constitutively-active ATPase activity in the charge-neutralized Asp824Asn (*12*) mutant implies that the salt-bridge formation between Lys791 and Asp824 drives the transport cycle forward from E2P. Another acidic side chain Asp942 (TM8) makes a salt bridge with Arg946 (3.3 Å), and is therefore likely deprotonated, despite being rather far from the K^+^-binding site. The deprotonation of Asp942 is substantiated in the MD simulations (Figure S6). Arg946 is replaced with Cys937 near the third Na^+^-binding site in Na^+^,K^+^-ATPase (*23*), and therefore the Asp942-Arg946 salt bridge observed in the H^+^,K^+^-ATPase structure can be predicted to be related to the pump’s electroneutral transport properties (*28*), although the function of the bridge is unclear in the absence of a high resolution E1 structure. Based on these observations, we conclude that Glu936 is protonated, and Asp824 and Asp942 are deprotonated. We further evaluated the protonation states of the K^+^-coordinating glutamate residues (E343, E795 and E820) by simulating various protonation state combinations (Figure 5).

**Figure 5.**
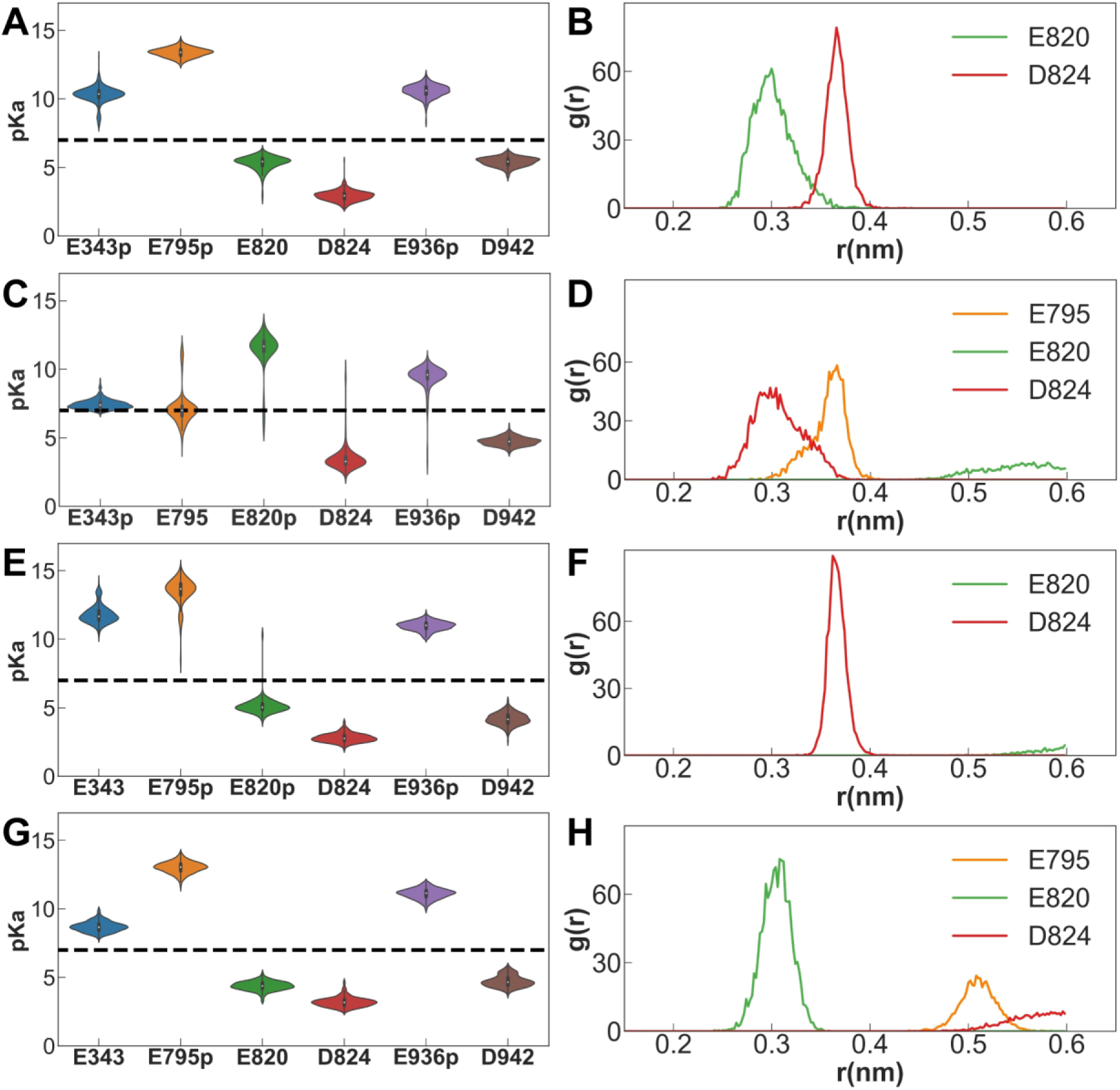
Molecular dynamics simulations of the K^+^-binding site. The *p*K_a_ distributions (**A, C, E** and **G**) over the last 50 ns of each simulation, and the radial distribution functions (RDF, **B, D, F** and **H**) between the ε-amino group of Lys791 and the center of mass of the indicated side chain carboxylate oxygen atoms of acidic residues during the MD simulations. The dashed horizontal line in the *p*K_a_ distributions indicates a *p*K_a_ of 7. The radial distribution function g(r) is a normalized histogram of distances between pairs of atom selections, averaged during the last 50 ns of each simulation.

The calculated average valence from each 50 ns (latter half of 100 ns simulation) × 3 copies of MD trajectories indicates that the protonation state expected from the crystal structure, namely, protonated Glu343, Glu795 and Glu936 (Figure 5A,B, for E343p/E795p/E936p) has a mean valence for K^+^ closest to the ideal value (1.07) in all examined simulation set-ups (Table S2). During the simulation, K^+^ was stably coordinated at the cation-binding site, and the root mean squared fluctuation (RMSF) of the ion remained below 1.0 Å in the likely protonation states. The calculated p*K*_a_ values of Glu343, Glu795 and Glu936 are above 7, and conversely those of Glu820, Asp824 and Asp942 are below 7, suggesting that the protonation status of E343p/E795p/E936p expected from the crystal structure is the most likely. The p*K*_a_ distribution plots from the different combinations of protonation states show that the estimated p*K*_a_ values are independent of the initial protonation assumption (note that the protonation state cannot change during the course of a single simulation). In fact, p*K*_a_ values of key acidic residues showed bimodal distribution in several other sets of simulations in which “wrong” protonation states were assumed (Figure 5, Figure S6). The sharp and unimodal p*K*_a_ distribution in E343p/E795p/E936p indicates a stably bound cation with stable conformations of the acidic side chains. The p*K*_a_ and RDF plots of E795p/E936p also show sharp distributions comparable to those of the likely protonation state of E343p/E795p/E936p. However, the pKa of Glu343 is well above 7 although it is kept de-protonated in the E795p/E936p simulation. Furthermore, the calculated valence for E795p/E936p is significantly higher (1.27) than that of E343p/E795p/E936p (Table S2). We therefore conclude that E343 is likely protonated, as expected from the crystal structure. Accordingly, Glu820 is the only deprotonated acidic residue that coordinates bound K^+^. However, the negative charge of Glu820 is neutralized by a stable salt bridge with Lys791 over the entire simulation period as seen in the RDF calculations, which is not observed at all in the simulations assuming Glu820 protonation (Figure 3, Figure S6).

Thus, K^+^ is not trapped by the negative charge in the cation binding site in spite of the crowded space and being surrounded by three glutamate residues. Like water molecules, oxygen atoms derived from either main chain carbonyl or protonated (E343 and E795) or charge-neutralized (E820) acidic side chains in the cation-binding site coordinate K^+^ by their lone pair electrons. Such binding, not driven by electrostatic interactions, will aid the release of K^+^ to the cytoplasmic solution with more than 100 mM K^+^. Stronger electrostatic interactions between K^+^ and the side chains would likely prevent the release of K^+^ even after a conformational change is exerted at the cation binding site in the E1 state.

## Discussion

Based on the crystal structure, and the protonation status suggested by MD simulation, the previously proposed transport model (*12*) need to be revised (Figure 6). In the K^+^-occluded E2-P transition state, a single K^+^ is observed at the cation-binding site of both wild-type enzyme and the Tyr799Trp mutant (Figure 2, Figure S2). Assuming electroneutral transport (*2,6*), one H^+^ must be exported in the prior luminal-open E2P state. Extensive hydrogen bonds and a salt bridge network centered on Glu820 in the luminal-open E2P state ensure that only one H^+^ is likely extruded from Glu820 regardless of the luminal pH (*12*). Accordingly, even when the luminal solution is at neutral pH, one H^+^ will remain bound to Glu343, which is suggested by both the crystal structures (Figure 3) and the molecular dynamics simulations (Figure 5).

**Figure 6.**
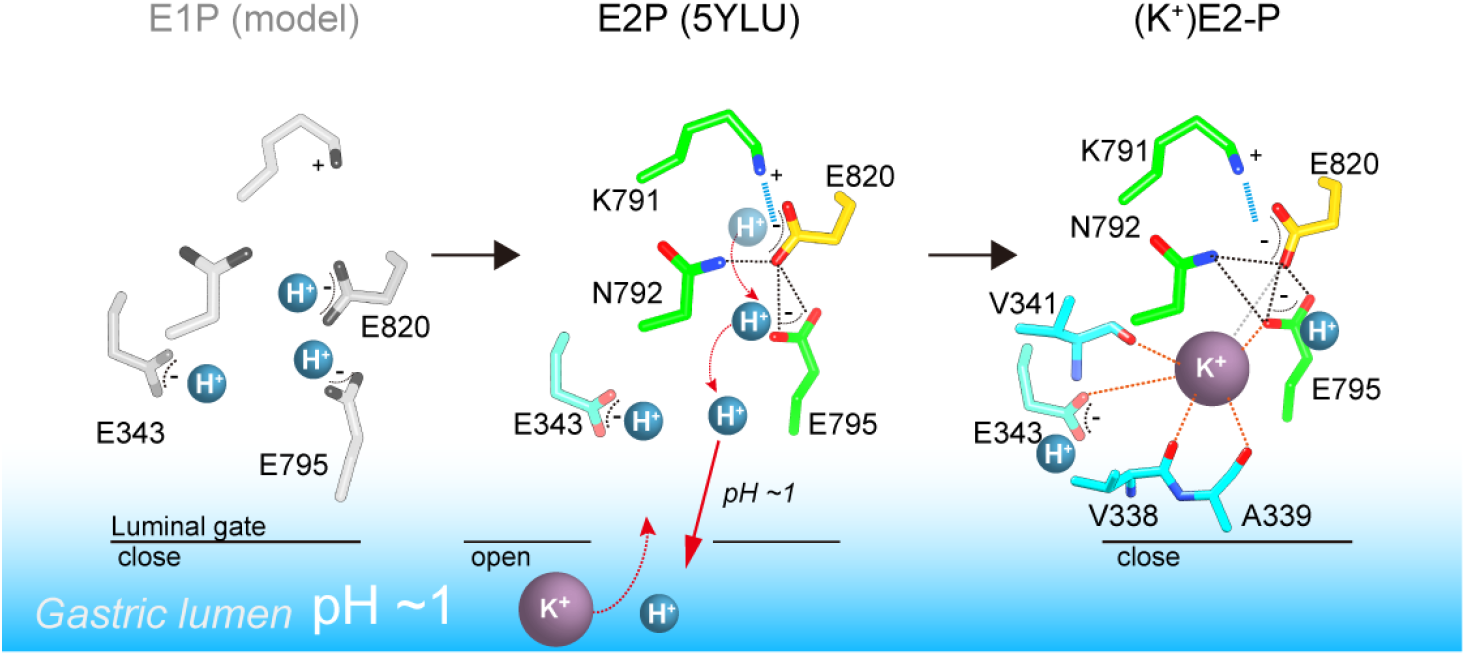
A model of one H^+^ and one K^+^ transport by H^+^,K^+^-ATPase. In the H^+^-occluded E1P state (left), all acidic residues are expected to be protonated. After a single H^+^ is extruded to the gastric luminal solution in the luminal-open E2P state (center), a single K^+^ is occluded in the following luminal-closed (K^+^)E2-P transition state (right). Dotted lines indicate hydrogen bonds (black), a salt bridge (blue), and K^+^-coordination by oxygen atoms (orange: ≤3 Å, grey >3 Å).

It is noteworthy that single K^+^ binding is driven by the Lys791-Glu820 interaction (Figure 3, 4), which also plays a key role in H^+^ extrusion into the acidic gastric solution (*12*). While the salt bridge is required for H^+^ extrusion, it also prevents binding of a second K^+^. Thus, H^+^,K^+^-ATPase seems to sacrifice 2H^+^/2K^+^ transport in order to achieve the energetically-challenging H^+^ transport of over a million-fold H^+^ gradient. The single K^+^ binding structure of H^+^,K^+^-ATPase therefore represents a remarkable example of how energetic barriers faced by membrane pumps are overcome in living systems. Evolutionary pressure selected single H^+^/K^+^ transport, rather than the more efficient double transport mode of 2H^+^/2K^+^, in order to achieve the thermodynamically challenging task of H^+^ uptake against a pH 1 solution in the stomach.

## Materials and Methods

### Protein expression and purification

Procedures for protein expression and purification are essentially the same as in a previous report (*12*). Briefly, wild-type (WT) or Tyr799Trp (YW) mutant of H^+^,K^+-^ATPase αβ-complex was expressed in the plasma membrane using baculovirus-mediated transduction of mammalian HEK293S GnT1^-^ cells (*29*). The harvested cells were broken up using a high-pressure emulsifier (Avestin), and membrane fractions collected. Membrane fractions were solubilized with 1% octaethylene glycol monododecyl ether (C_12_E_8_, Nikko Chemical) with 40 mM MES/Tris (pH 6.5), 10% glycerol, 5 mM dithiothreitol in the presence of 50 mM CH_3_COORb, 10 mM MgCl_2_, 10 mM NaF for WT(Rb^+^)E2-MgF, or 200 mM KCl, 10 mM MgCl_2_, 10 mM NaF for YW(K^+^)E2-MgF, or 200 mM RbCl, 10 mM MgCl_2_, 10 mM NaF for YW(Rb^+^)E2-MgF or 200 mM RbCl, 1 mM MgCl_2_, 1 mM AlCl_3_, 4 mM NaF for YW(Rb^+^)E2-AlF, on ice for 20 min. Proteins were affinity purified by anti-Flag M2 affinity resin (Sigma-Aldrich), which followed digestion of affinity tag and deglycosilation by TEV protease and MBP-fusion endoglycosidase (New England Biolabs), respectively, at 4 °C overnight. Samples were further purified by a size-exclusion column chromatograph using a Superose6 Increase column (GE Healthcare). Peak fractions were collected and concentrated to 10 mg/ml. The concentrated H^+^,K^+^-ATPase samples were added to the glass tubes in which a layer of dried dioleoyl phosphatidylcholine had formed, in a lipid-to-protein ratio of 0.1-0.4, and incubated overnight at 4 °C in a shaker mixer operated at 120 rpm. After removing the insoluble materials by ultracentrifugation, the lipidated samples were used for the crystallization.

### Crystallization

Initial screen was performed using a K^+^ salt-containing matrix called K^+^night screen, which was developed based on the King screen (*30*). Crystals were obtained by vapor diffusion at 20 °C. For the YW mutant, a 5-mg/ml purified, lipidated protein sample was mixed with reservoir solution containing 10% glycerol, 20% PEG2000MME, 3% methylpentanediol, and 5 mM β-mercaptoethanol in the presence of 0.4 M KCl for YW(K^+^)E2-MgF state, or 0.4 M RbCl for YW(Rb^+^)E2-MgF and YW(Rb^+^)E2-AlF states. For the WT enzyme, reservoir solution containing 10% glycerol, 15% PEG6000, 0.1 M CH_3_COORb, 6% methylpentanediol and 5 mM β-mercaptoethanol was used. Crystals were flash frozen in liquid nitrogen.

### Structural determination and analysis

Diffraction data were collected at the SPring-8 beamline BL32XU and BL41XU, and processed using XDS. Structure factors were subjected to anisotropy correction using the UCLA MBI Diffraction Anisotropy server (*31*) (http://services.mbi.ucla.edu/anisoscale/). The structure of YW(Rb^+^)E2-AlF was determined by molecular replacement with PHASER, using an atomic model of H^+^,K^+^-ATPase in the SCH28080-bound E2BeF state (pdb ID: 5YLV) as a search model. Coot (*32*) was used for cycles of iterative model building and Refmac5 and Phenix (*33*) were used for refinement. Other structures described in this paper were determined by molecular replacement using an atomic model of YW(Rb^+^)E2-AlF_4_^-^ state as a search model. Rubidium and potassium ions were identified in anomalous difference Fourier maps calculated using data collected at wavelengths of 0.8147 and 1.700 Å, respectively. The YW(K^+^)E2-MgF_4_^2-^, YW(Rb^+^)E2-MgF_4_^2-^, YW(Rb^+^)E2-AlF_4_^-^ and WT(Rb^+^)E2-MgF_4_^2-^ models contained 98.2/1.8/0.0 %, 98.3/1.7/0.0 %, 98.2/1.8/0.0 % and 91.1/8.8/0.1% in the favored, allowed, and outlier regions of the Ramachandran plot, respectively.

### Activity assay using recombinant proteins

The wild-type or mutant α-subunit was co-expressed with the wild-type β-subunit using the BacMam system as described above, and broken membrane fractions were collected. H^+^,K^+^-ATPase activity was measured as described previously (*34*). Briefly, permeabilized membrane fractions (wild-type or mutant) were suspended in buffer comprising 40 mM PIPES/Tris (pH 7.0), 2 mM MgCl_2_, 2 mM ATP, and 0-50 mM KCl in the presence of three different concentrations of vonoprazan, or their absence, in the 96-well plates. Reactions were initiated by incubating the fractions at 37 °C using a thermal cycler, and maintained for 1 to 5 h depending on their activity. Reactions were terminated, and the amount of released inorganic phosphate was determined colorimetrically using a microplate reader (TECAN).

### Molecular dynamics simulations

All simulations were performed using GROMACS (v-2016.3) (*35–39*). The CHARMM36 force field (v-July 2017) (*40–43*) was used to model all the components of the system. A symmetric lipid bilayer containing ~500 DOPC lipids was generated using CHARMM-GUI (*44,45*). An atomic model of H^+^, K^+^-ATPase YW(K^+^)E2-MgF_4_^2-^ was placed in the bilayer using the OPM (*46*) server. All the components of experimentally derived structure were kept intact in the system except detergent and MgF_4_^2-^ which was replaced with a PO_4_^3-^ molecule. Systems were hydrated with ~81000 TIP3P water molecules and ionized with 150 mM NaCl.

Neighbor search for non-bonded interactions was performed within a cutoff of 12 Å and revised after every 20 steps. Van der Waals forces were smoothly switched off between 10 Å and 12 Å with a force-switch function. The particle-mesh Ewald (PME) (*47,48*) method was used to calculate electrostatic interactions. Initially, an energy minimization was performed using the gradient descent method to remove steric clashes. Then, systems were equilibrated for a total of 25-30 ns in three steps. In the first step, heavy atoms of the protein including the potassium ions were restrained with a force constant of 1000 kJ/mol·nm^2^, which was reduced to zero in two further equilibration steps. Finally, a production run was performed for 100 ns without restraints. Periodic boundary conditions were incorporated. All acidic residues pointing towards the stomach lumen were kept protonated.

Overall, 18 protonation states with either 2, 3 or 4 acidic protonated residues were simulated. For some protonation states that were deemed more likely, based on the analysis, we launched two further copies of the simulations with different initial velocity distributions. The results from all simulation copies are nearly identical. A time-step of 1 fs was used for the simulations. The temperature of the systems was stabilized at the 310 K by integrating the Nosé-Hoover thermostat (*49,50*) during the production runs. The Parrinello-Rahman barostat (*51*) along with a semiisotropic pressure coupling scheme were used to maintain the pressure at 1 bar. All bonds containing hydrogen were constrained using The Linear Constraint Solver algorithm (LINCS) (*52*). PROPKA (v-3.1) (*53–55*) was used to measure p*K*_a_ dynamically at every 100 ps. Only the last 50 ns were considered for the data analysis. GROMACS and python scripts were used for the various analyses used in the manuscript. Snapshots were generated using Visual Molecular Dynamics (VMD) (*56*).

## Acknowledgments

We thank M. Taniguchi for the technical assistance; D. McIntosh for improving the manuscript; C. Toyoshima for discussions. The synchrotron radiation experiments were performed at BL32XU and BL41XU in SPring-8 with the approval of the Japan Synchrotron Radiation Research Institute (JASRI Proposal numbers: 2017B2701 and 2018B2703). We thank the beamline staff for their facilities and support. The simulations were carried out on the Danish e-Infrastructure Cooperation (DeiC) National HPC Center, ABACUS 2.0 at the University of Southern Denmark, SDU as well as on computing resources on the Swiss cluster Piz Daint as part of the PRACE grant number 2016153468.

## Funding

This work was supported by JSPS KAKENHI Grant Number 17H03653, JST CREST (JPMJCR14M4), AMED BINDS and Takeda Science Foundation (to K.A.); the Japan New Energy and Industrial Technology Development Organization (NEDO), and the Japan Agency for Medical Research and Development (AMED) (to Y.F.). H.K. is supported by Lundbeckfonden. V.D. is supported by Lundbeckfonden and the e-Science Center SDU.

## Author contributions

K.A. and Y.F. designed the study. K.Y., H.N. and K.A. expressed the proteins. K.Y. and K.A. purified and crystallized the proteins. K.A. performed the biochemical analysis. K.Y. and K.A. collected the X-ray diffraction data. K.I. and K.A. analyzed the structure. V.D. and H.K. designed the MD simulations. V.D. performed and analyzed the simulations. K.A. interpreted the structure and K.A. and H.K. wrote the manuscript, with agreement of all authors.

## Competing interests

Y.F. is a director of CeSPIA Inc.

## Data and materials availability

Atomic coordination and structure factors for the structures reported in this work were deposited in the Protein Data Bank under accession number XXXX for YW(K^+^)E2-MgF, YYYY for YW(Rb^+^)E2-MgF, ZZZZ for YW(Rb^+^)E2-AlF and WWWW for WT(Rb^+^)E2-MgF.

## Supplementary Materials

Figures S1-S6

Tables S1-S2

Movies S1-S2

