## Supplemental figures and tables for "A single K^+^-binding site in the crystal structure of the gastric proton pump"

**This PDF file includes:**

Figures S1 to S6
Tables S1 to S2
Captions for Movies S1 to S2

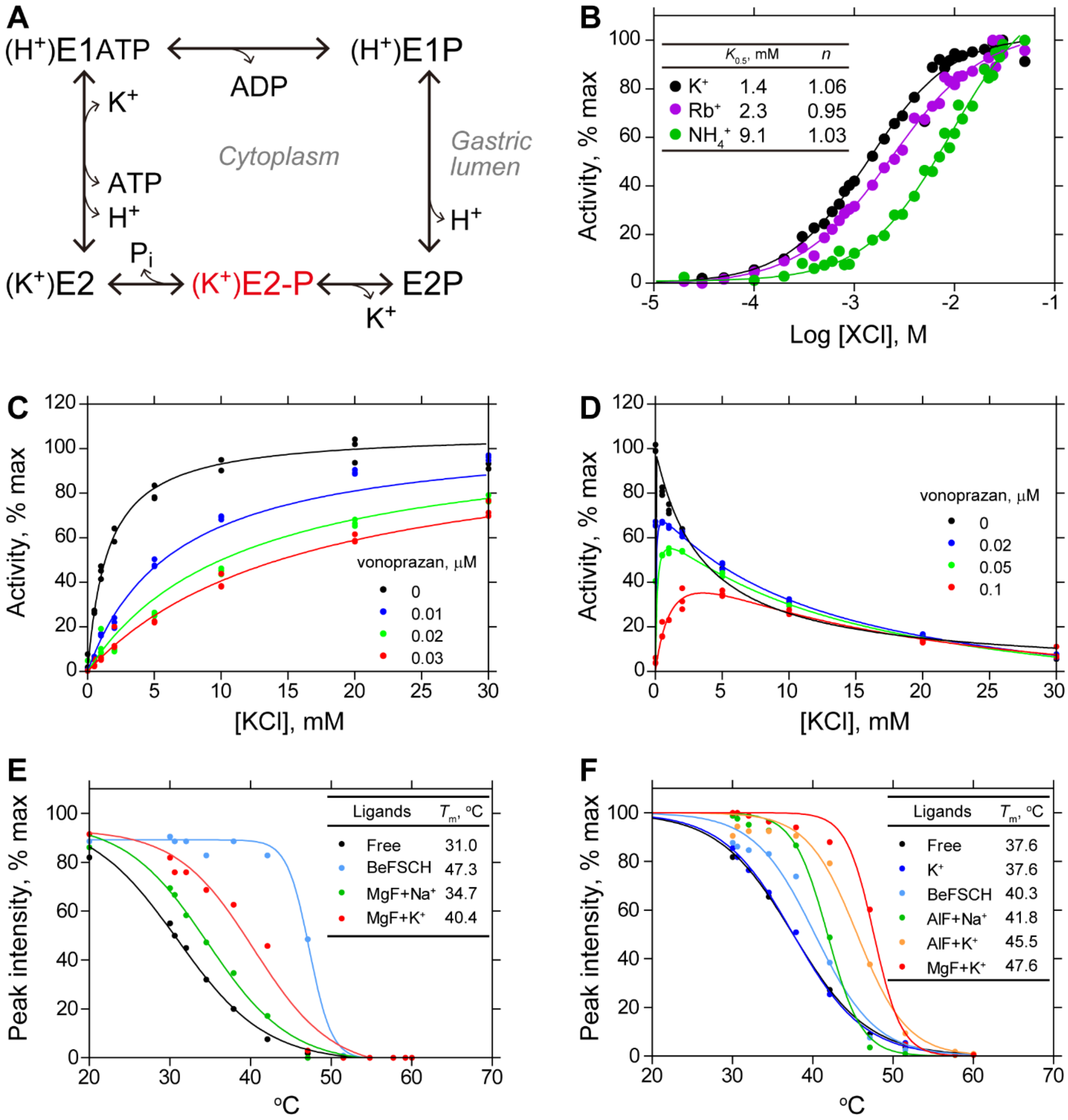

**Figure S1. Characterization of Tyr799Trp mutant of $H^+$,$K^+$-ATPase.**

**A**, Post-Albers type reaction scheme for $H^+$,$K^+$-ATPase. $K^+$-occluded E2-P transition state is highlighted in red. **B**, ATPase activities of the wild-type enzyme with indicated cations, showing Hill coefficient (*n*) close to 1. $K^+$-dependent ATPase activity of wild-type (**C**) or Try799Trp mutant (**D**), in the absence or presence of indicated concentrations of $K^+$-competitive inhibitor vonoprazan. Thermal stability of wild-type (**E**) or Try799Trp (**F**) in the presence of indicated ligands, evaluated by fluorescence size-exclusion chromatography.

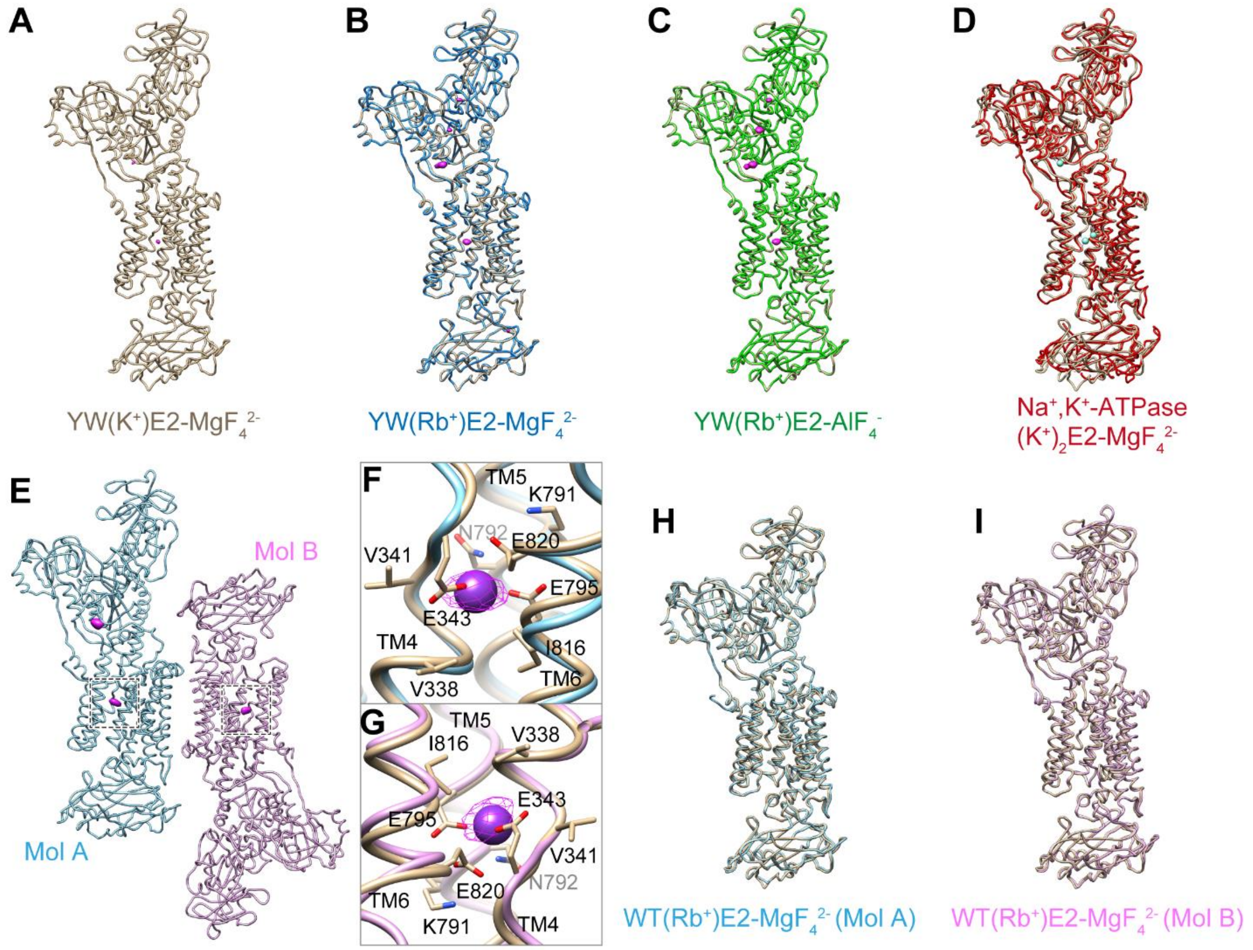

**Figure S2. Comparison of the molecular conformation for $K^+$ or $Rb^+$-occluded E2-P transition state of the $H^+$,$K^+$-ATPase.**

Comparison of the molecular conformation of the $H^+$,$K^+$-ATPase YW($K^+$)E2-$MgF_4^{2-}$ (**A**, wheat) and YW($Rb^+$)E2-$MgF_4^{2-}$ (**B**, blue), YW($Rb^+$)E2-$AlF_4^-$ (**C**, green), or $Na^+$,$K^+$-ATPase (2$K^+$)E2-$MgF_4^{2-}$ (**D**, red, 2zxe). Asymmetric unit of wild-type $H^+$,$K^+$-ATPase in the ($Rb^+$)E2-$MgF_4^{2-}$ state contains two αβ-complexes (**E**). Close-up of the cation-binding site in Mol A (**F**, light blue) and Mol B (**G**, pink) indicated dotted boxes in **e**. Comparison of the molecular conformation of WT($Rb^+$)E2-$MgF_4^{2-}$ and YW($K^+$)E2-$MgF_4^{2-}$ (**H**,**I**). Anomalous difference Fourier maps of indicated crystal structures are shown as magenta in **A**-**G** with contour level of 4σ for $K^+$(**A**); 8σ for $Rb^+$ (**B**-**D**); 4σ for $Rb^+$ (**E**-**G**).

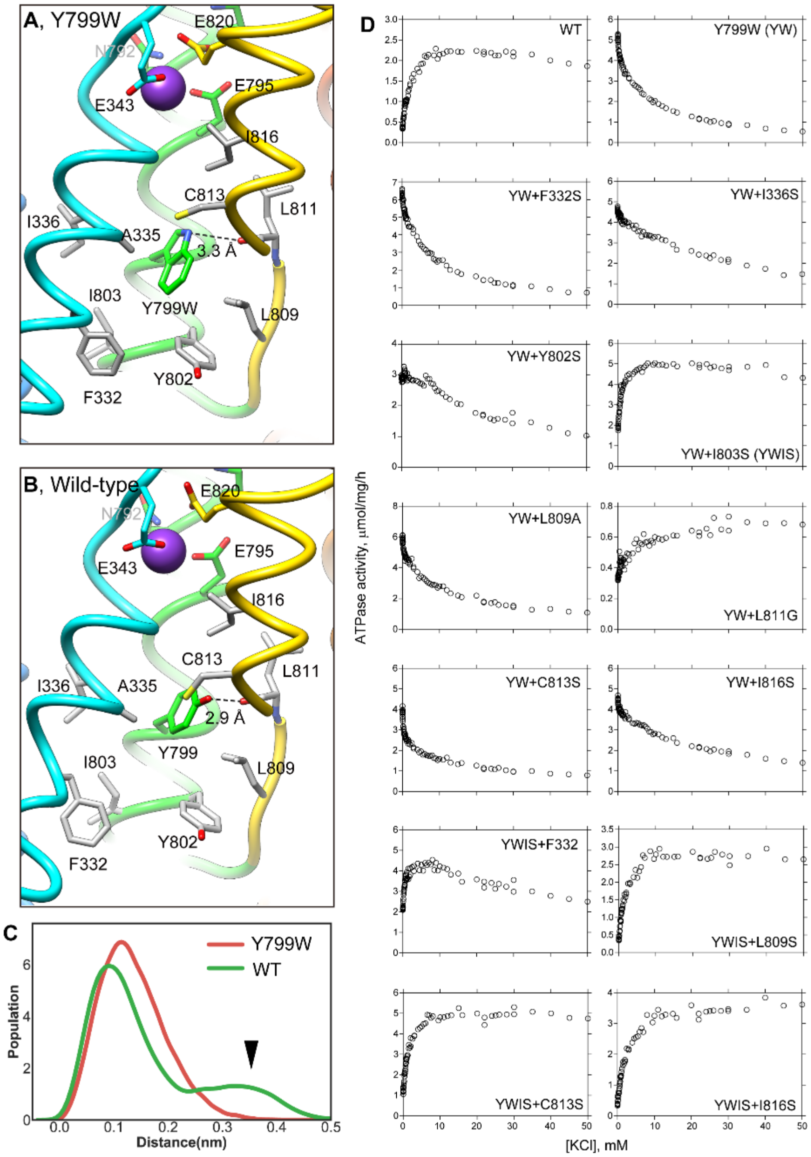

**Figure S3. K⁺-dependence on ATPase activities of Y799W and mutant of surrounding hydrophobic amino acids.**

Hydrophobic interactions observed in the luminal gate of Y799W (**A**) and WT (**B**). Hydrophobic residues likely contribute to the luminal gate stability are shown as grey sticks. (**C**) Distribution of the root-mean square deviations (RMSD) for NH group of Trp residue in Y799W (red line, mean RMSD is 1.23 ± 0.04) and OH group of Tyr799 in WT (green line, 1.59 ± 0.27). The arrowhead points to a population of large RMSD, indicating higher fluctuations of the OH group in WT, compared to the NH group in the Tyr799Trp mutant (see also Movie S1). (**D**) $K^+$-dependence of vonoprazan-sensitive ATPase activities of the indicated mutants. Data plotted had background values in the presence of 10 μM vonoprazan subtracted. Individual data from a 96-well plate at different concentrations of $K^+$ were plotted, and representative results from more than three independent measurements for each mutant are shown in the figure.

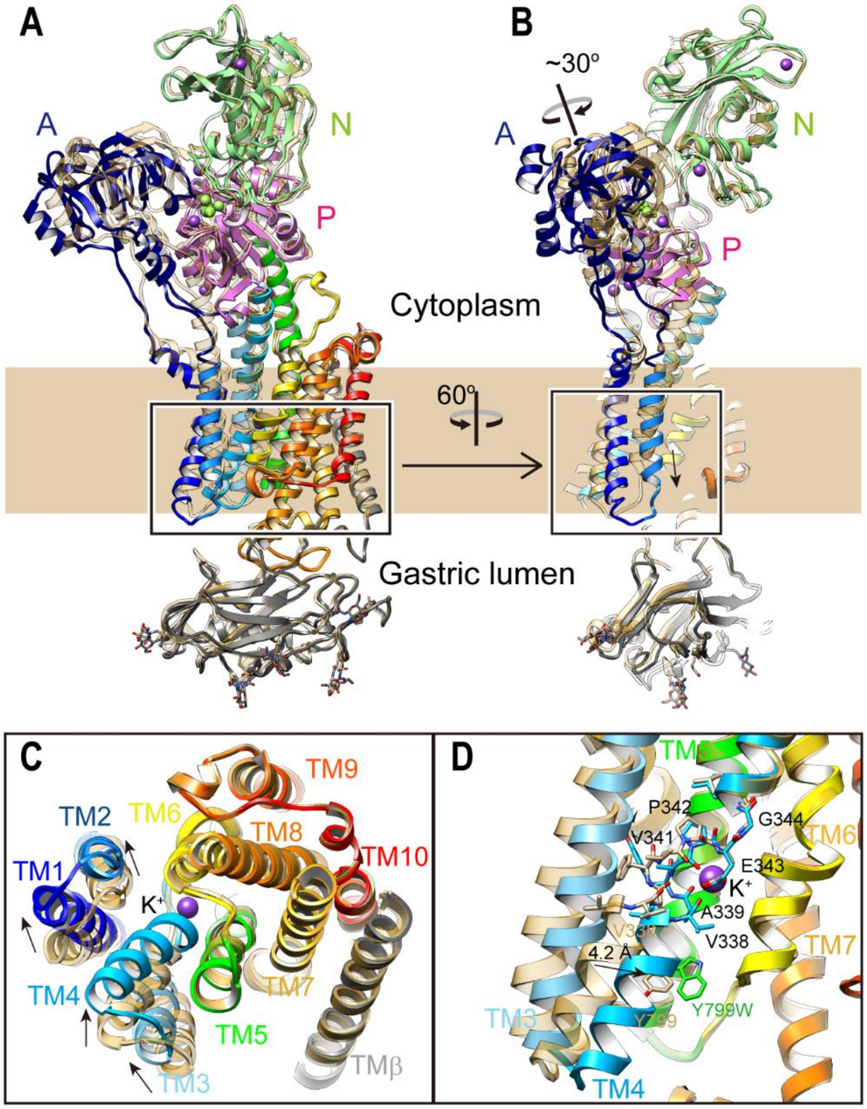

**Figure S4. Conformational change upon $K^+$-occlusion.**

Luminal-open E2P state [pdb code: 5YLU, (von)E2$BeF_3^-$, wheat ribbons] is superimposed on the YW($K^+$)E2-$MgF_4^{2-}$ structure (color codes as in Fig. 1), showing the conformational change in the whole complex (**A**,**B**). **C**, Rearrangement of TM helices upon $K^+$-binding. TM helices indicated in the box in **A** were viewed from the luminal side of the membrane. **D**, Displacement of TM4 upon luminal gate closure. Figure shows an enlargement of the indicated a box in **B**, viewed from parallel to the membrane normal. The unwound region of TM4 is shown as stick representations. TM1 and TM2 were omitted for clarity. Arrows indicate the displacement from luminal-open E2P to luminal-closed ($K^+$)E2-P states.

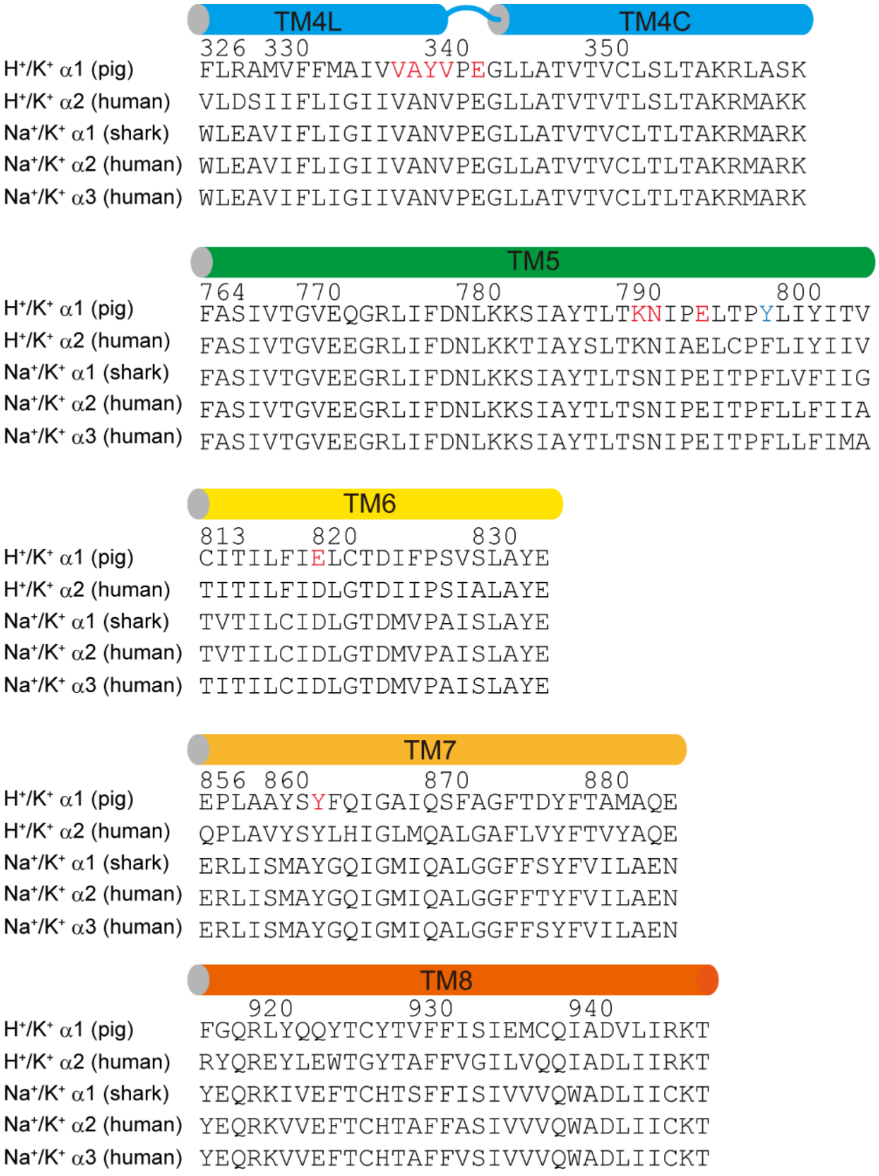

**Figure S5. Sequence alignment of TM helices of $H^+$,$K^+$- and $Na^+$,$K^+$-ATPases.**

Sequence alignment of the indicated transmembrane helices among related P2C-type ATPases. The amino acids explicitly discussed in the manuscript are highlighted in red. Tyr799, which is changed to Trp in the high-resolution crystal structure of $H^+$,$K^+$-ATPase is indicated in blue.

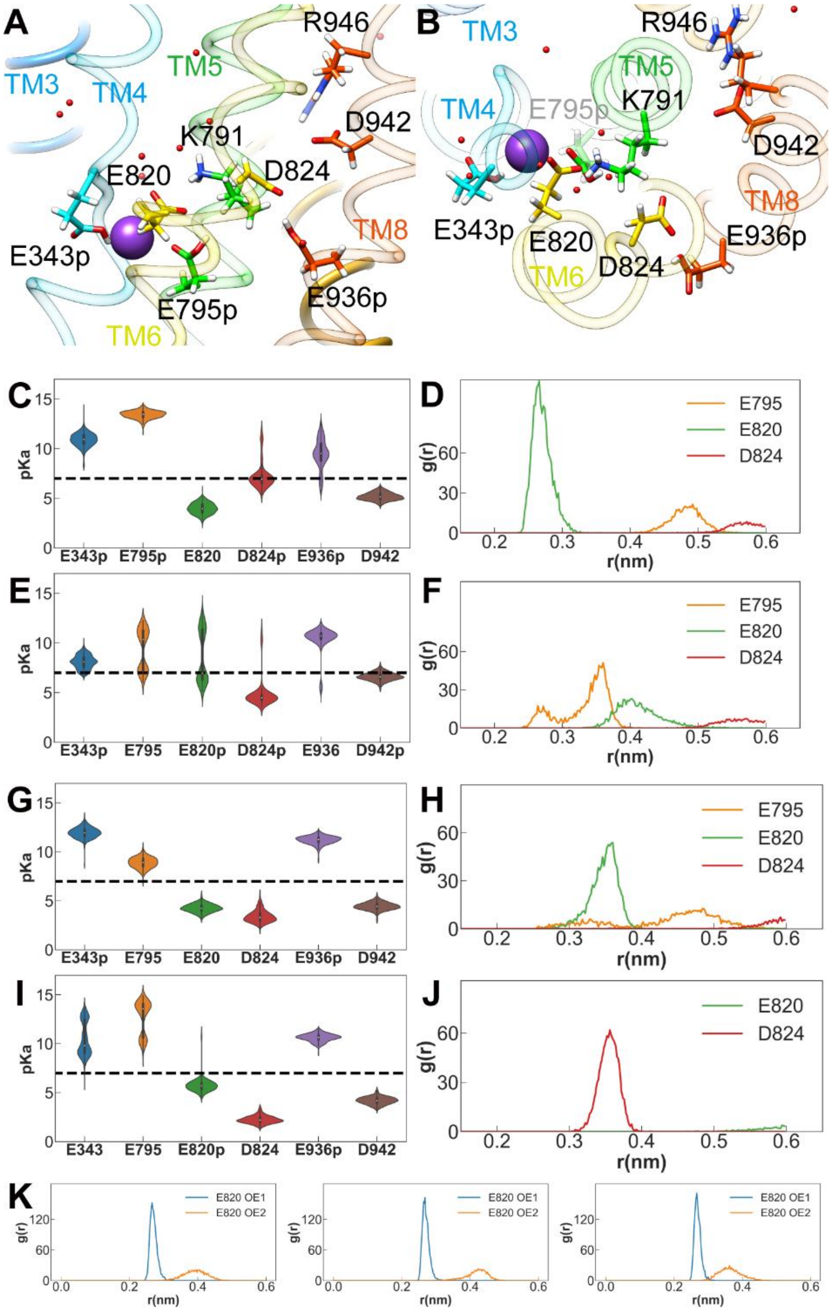

**Figure S6. Molecular dynamics simulation of $K^+$-occluded $H^+$,$K^+$-ATPase.**

**A**,**B**, An experimental set-up of the cation-binding site for the MD simulation with protonated Glu343, Glu795 and Glu936, showing the valence closest to the ideal value among evaluated simulations (see Table S2). Several acidic side chains near the cation-binding site, and Lys791 are only shown as sticks with their hydrogen atoms. Calculated p$K_a$ values (**C**,**E**,**G**,**I**) and RDFs for Lys791 and indicated side chains (**D**,**F**,**H**,**J**) for each indicated simulation. **H**, RDF plots of Lys791 Nε and Glu820 Oε1 or Oε2 in the three independent simulations of the most likely protonation state (Fig. 3, E343p/E795p/E936p), showing the stable salt bridge between them. Figures were prepared as in Fig. 3. See Table S2 for the valence in each simulation.

| | YW($K^+$)E2-$MgF_4^{2-}$ | YW($Rb^+$)E2-$MgF_4^{2-}$ | YW($Rb^+$)E2-$AlF_4^-$ | WT($Rb^+$)E2-$MgF_4^{2-}$ |
|---|---|---|---|---|
| **Data collection** | | | | |
| Resolution (Å) † | 2.7×2.8×2.5 (2.6 - 2.5)‡ | 2.8×2.8×2.6 (2.7 - 2.6) | 2.8×3.0×2.5 (2.6 - 2.5) | 5.1×5.1×4.3 (4.5 – 4.3) |
| Space group | $P\,3_1\,2\,1$ | $P\,3_1\,2\,1$ | $P\,3_1\,2\,1$ | $C$ 1 2 1 |
| Cell dimensions | | | | |
| *a, b, c* (Å) | 103.37, 103.37, 370.01 | 103.38,103.38,369.86 | 103.23,103.23,369.44 | 191.51,106.43,250.96 |
| α, β, γ (˚) | 90, 90, 120 | 90, 90, 120 | 90, 90, 120 | 90, 107.79, 90 |
| $R_{merge}$ | 0.067 (3.685) | 0.080 (2.248) | 0.106 (3.499) | 0.152 (1.359) |
| $R_{pim}$ | 0.022 (1.179) | 0.030 (0.855) | 0.040 (1.295) | 0.065 (0.544) |
| $I/\sigma I$ | 20.18 (0.93) | 16.19 (1.16) | 11.64 (0.77) | 7.19 (0.92) |
| *C/C1/2* | 0.99 (0.59) | 0.99 (0.70) | 0.99 (0.60) | 0.99 (0.75) |
| Completeness (%) | 85.65 (26.79) | 88.93 (36.11) | 76.00 (18.98) | 94.94 (53.46) |
| Redundancy | 10.2 (10.6) | 8.0 (7.8) | 8.0 (8.2) | 6.6 (7.2) |
| **Refinement** | | | | |
| Resolution (Å) | 48 – 2.5 (2.6 - 2.5) | 48 – 2.6 (2.7 - 2.6) | 50 – 2.5 (2.6 - 2.5) | 48 – 4.3 (4.5 – 4.3) |
| No. of reflections | 69037 (2135) | 63831 (2533) | 61013 (1502) | 31390 (1747) |
| $R_{work}$/ $R_{free}$ (%) | 20.1/25.7 (37.3/48.0) | 20.6/26.2 (35.1/43.5) | 21.5/27.4 (32.7/42.6) | 26.3/33.9 (33.4/41.2) |
| Wilson B-factor | 60.74 | 52.60 | 48.43 | 158.52 |
| No. of atoms | 10513 | 10485 | 10498 | 19823 |
| Protein | 9736 | 9762 | 9763 | 19623 |
| Ligands | 362 | 362 | 364 | 200 |
| Average B-factor | 74.40 | 62.68 | 60.03 | 244.07 |
| Protein (Å$^2$) | 72.62 | 60.98 | 57.83 | 243.40 |
| Ligands (Å$^2$) | 122.80 | 114.14 | 103.16 | 310.29 |
| R.m.s deviations | | | | |
| Bond lengths (Å) | 0.010 | 0.009 | 0.010 | 0.005 |
| Bond angles (˚) | 1.36 | 1.51 | 1.39 | 1.18 |

**Table S1. Data collection and refinement statistics.**

†The diffraction data are anisotropic. The resolution limits given are for the *a**, *b** and *c** axes, respectively.

‡Statistics for the highest-resolution shell are shown in parentheses.

| | 2-proton states | | | 3-proton states | | | 4-proton states | |
|---|---|---|---|---|---|---|---|---|
| E343 | $H^+$ | - | - | $H^+$ | $H^+$ | - | $H^+$ | $H^+$ |
| E795 | - | $H^+$ | - | $H^+$ | - | $H^+$ | $H^+$ | - |
| E820 | - | - | $H^+$ | - | $H^+$ | $H^+$ | - | $H^+$ |
| D824 | - | - | - | - | - | - | $H^+$ | $H^+$ |
| E936 | $H^+$ | $H^+$ | $H^+$ | $H^+$ | $H^+$ | $H^+$ | $H^+$ | - |
| D942 | - | - | - | - | - | - | - | $H^+$ |
| | Valence† (Mean ± SEM) | | | | | | | |
| V338 O | 0.077±0.001 | 0.239±0.004 | 0.282±0.002 | 0.042±0.002 | 0.116±0.004 | 0.268±0.001 | 0.111±0.003 | 0.217±0.001 |
| A339 O | 0.132±0.001 | 0.098±0.002 | 0.005±0.000 | 0.171±0.004 | 0.235±0.002 | 0.128±0.001 | 0.203±0.003 | 0.248±0.001 |
| V341 O | 0.242±0.001 | 0.278±0.005 | 0.224±0.001 | 0.239±0.003 | 0.269±0.001 | 0.283±0.002 | 0.295±0.003 | 0.29±0.001 |
| E343 Oε1 | 0.21±0.001 | 0.169±0.004 | 0.257±0.002 | 0.046±0.002 | - | 0.26±0.002 | 0.068±0.003 | - |
| E343 Oε2 | 0.005±0.000 | 0.152±0.003 | 0.284±0.002 | 0.002±0.000 | - | 0.206±0.002 | 0.002±0.000 | - |
| E795 Oε1 | 0.278±0.003 | 0.121±0.002 | 0.291±0.002 | 0.267±0.003 | 0.315±0.002 | 0.107±0.001 | 0.277±0.003 | 0.239±0.003 |
| E795 Oε2 | 0.134±0.003 | -‡ | 0.272±0.002 | - | 0.053±0.002 | 0.001±0.000 | - | 0.239±0.003 |
| E820 Oε1 | 0.200±0.002 | 0.017±0.001 | 0.001±0.000 | 0.015±0.000 | 0.061±0.001 | 0.001±0.000 | 0.138±0.005 | 0.155±0.001 |
| E820 Oε2 | 0.109±0.002 | 0.195±0.004 | 0.010±0.000 | 0.291±0.006 | 0.089±0.001 | 0.028±0.000 | 0.118±0.005 | 0.066±0.001 |
| total | 1.387 | 1.269 | 1.626 | 1.073¶ | 1.138 | 1.282 | 1.212 | 1.454 |

**Table S2. Mean valences calculated in the $K^+$-binding site of $H^+$,$K^+$-ATPase assuming protonation for the acidic residues as evaluated by MD simulations.**

†Ideal value for $K^+$ is 1.00. ‡Only oxygen atoms within 4 Å from bound $K^+$ were included for the valence calculation. ¶The most likely protonation state of E343p/E795p/E936p shows a total valence closest to the ideal value amongst all simulations evaluated.

**Movie S1. MD simulation of Tyr799Trp mutant and the wild-type enzymes.**
A movie shows the luminal gate and $K^+$-binding site structure of 100 ns MD simulations for Tyr799Trp mutant (left) and the wild-type enzyme.

**Movie S2. $K^+$-binding site.**
Detailed structure of the $K^+$-binding site in YW($K^+$)E2-$MgF_4^{2-}$ state, viewed as in Fig. 2A.
